# Ensemble Machine Learning Modeling for the Prediction of Artemisinin Resistance in Malaria

**DOI:** 10.1101/856922

**Authors:** Colby T. Ford, Daniel Janies

## Abstract

Antiparasitic resistance in malaria is a growing concern affecting many areas of the eastern world. Since the emergence of artemisinin resistance in the late 2000s in Cambodia, research into the underlying mechanisms has been underway.

The 2019 Malaria Dream Challenge posited the task of developing computational models that address important problems in advancing the fight against malaria. The first goal was to accurately predict Artemisinin drug resistance levels of *Plasmodium falciparum* isolates, as quantified by the IC_50_. The second goal was to predict the parasite clearance rate of malaria parasite isolates based on *in vitro* transcriptional profiles.

In this work, we develop machine learning models using novel methods for transforming isolate data and handling the tens of thousands of variables that result from these data transformation exercises. This is demonstrated by using massively parallel processing of the data vectorization for use in scalable machine learning. In addition, we show the utility of ensemble machine learning modeling for highly effective predictions of both goals of this challenge. This is demonstrated by the use of multiple machine learning algorithms combined with various scaling and normalization preprocessing steps. Then, using a voting ensemble, multiple models are combined to generate a final model prediction.

## Introduction

Malaria is a serious disease caused by parasites belonging to the genus *Plasmodium* which are vectored by *Anopheles* mosquitoes in the genus. The World Health Organization (WHO) reports that there were 219 million cases of malaria in 2017 across 87 countries^1^. *Plasmodium falciparum* poses one of greatest health threats in the eastern world, being responsible for 62.8% of malaria cases in southeast Asia in 2017^1^.

Artemisinin-based therapies are among the best treatment options for malaria caused by *P. falciparum*^2^. However, emergence of artemisinin resistance in Thailand and Cambodia in 2007 has been cause for research^3^. While there are polymorphisms in the kelch domain–carrying protein K13 in *P. falciparum* that are known to be associated with artemisinin resistance, the underlying molecular mechanism that confers resistance remain unknown^4^. The established pharmacodynamics benchmark for *P. falciparum* sensitivity to artemisinin-based therapy is the parasite clearance rate^5, 6^. Resistance to artemisinin-based therapy is considered to be present with a parasite clearance rate greater than 5 hours^7^. By understanding the genetic factors that affect resistance in malaria, targeted development can occur in an effort to abate further resistance or infections of resistance strains.

## Prediction of Artemisinin IC_50_

First, we created a machine learning model to predict the IC_50_ of malaria parasites based on transcription profiles of experimentally-tested isolates. IC_50_, also known as the half maximal inhibitory concentration, is the drug concentration at which 50% of parasites die. This value indicates a population of parasites’ ability to withstand various doses of anti-malarial drugs, such as Artemisinin.

### Materials and Methods

The training data, from Turnbull, et al.^8^, consists of gene expression data of 5,540 genes of 30 isolates from the malaria parasite, *Plasmodium falciparum*. For each malaria parasite isolate, transcription data was collected at two time points [6 hours post invasion (hpi) and 24 hpi], with and without treatment of dihydroartemisinin (the metabolically active form of artemisinin), each with a biological replicate. This yields a total of at eight data points for each isolate. The initial form of the training dataset contains 272 rows and 5,546 columns, as shown in Table 1.

**Table 1.** Initial IC_50_ model training data format.

| Sample_Name | Isolate | Timepoint | Treatment | BioRep | Gene<sub>1</sub> | ... | Gene<sub>5540</sub> | DHA_IC50 |
| --- | --- | --- | --- | --- | --- | --- | --- | --- |
| isolate_01.24HR.DHA.BRep1 | isolate_01 | 24HR | DHA | BRep1 | 0.008286 | ... | -2.48653 | 2.177 |
| isolate_01.24HR.DHA.BRep2 | isolate_01 | 24HR | DHA | BRep2 | -0.87203 | ... | -1.79457 | 2.177 |
| isolate_01.24HR.UT.BRep1 | isolate_01 | 24HR | UT | BRep1 | 0.03948 | ... | -2.49517 | 2.177 |
| isolate_01.24HR.UT.BRep2 | isolate_01 | 24HR | UT | BRep2 | 0.125177 | ... | -1.73531 | 2.177 |
| isolate_01.6HR.DHA.BRep1 | isolate_01 | 6HR | DHA | BRep1 | 1.354956 | ... | -0.82169 | 2.177 |
| isolate_01.6HR.DHA.BRep2 | isolate_01 | 6HR | DHA | BRep2 | -0.21807 | ... | -1.61839 | 2.177 |
| isolate_01.6HR.UT.BRep1 | isolate_01 | 6HR | UT | BRep1 | 1.31135 | ... | -2.62262 | 2.177 |
| isolate_01.6HR.UT.BRep2 | isolate_01 | 6HR | UT | BRep2 | 0.997722 | ... | -2.24719 | 2.177 |
| ... | ... | ... | ... | ... | ... | ... | ... | ... |
| isolate_30.6HR.UT.BRep2 | isolate_30 | 6HR | UT | BRep2 | -0.26639 | ... | -1.72273 | 1.363 |

The transcription data was collected as described in Table 2. The transcription data set consists of transcription values for 100 *MAL* genes (SNARE protein-coding genes^9^) followed by 5,440 *PF3D7* genes (circumsporozoite protein-coding genes^10^). The *MAL* genes are 92 non-coding RNAs while the *PF3D7* genes are protein coding genes. The feature to predict is *DH*A_*IC*50.

**Table 2.** IC_50_ training data information. (Adapted from Turnbull et al., (2017) PLoS One^8^)

|  | Training Set |
| --- | --- |
| Array | Bozdech |
| Platform | Printed |
| Plexes | 1 |
| Unique Probes | 10159 |
| Range of Probes per Exon | N/A |
| Average Probes per Gene | 2 |
| Genes Represented | 5363 |
| Transcript Isoform Profiling | No |
| ncRNAs | No |
| Channel Detection Method | Two Color |
| Scanner | PowerScanner |
| Data Extraction | GenePix Pro |

### Data Preparation

We used Apache Spark^11^, to pivot the dataset such that each isolate was its own row and each of the transcription values for each gene and attributes (i.e. timepoint, treatment, biological replicate) combination was its own column. This exercise transformed the training dataset from 272 rows and 5,546 columns to 30 rows and 44,343 columns, as shown in Table 3.

**Table 3.** Post-transformation format of the IC_50_ model training data.

| Isolate | DHA_IC50 | hr24_trDHA_br1_Gene<sub>1</sub> | hr24_trDHA_br2_Gene<sub>1</sub> | ... | hr6_trUT_br2_Gene<sub>5540</sub> |
| --- | --- | --- | --- | --- | --- |
| isolate_01 | 2.177 | 0.008286 | -0.87203 | ... | -2.24719 |
| ... | ... | ... | ... | ... | ... |
| isolate_30 | 1.363 | 0.195032 | 0.031504 | ... | -1.72273 |

We completed this pivot by slicing the data by each of the eight combinations of timepoint, treatment, and biological replicate, dynamically renaming the variables (genes) for each slice, and then joining all eight slices back together.

Example code shown below in the section labeled code 1. By using the massively parallel architecture of Spark, this transformation can be completed in a minimal amount of time on a relatively small cluster environment (e.g., <10 minutes using a 8-worker/36-core cluster with PySpark on Apache Spark 2.4.3).

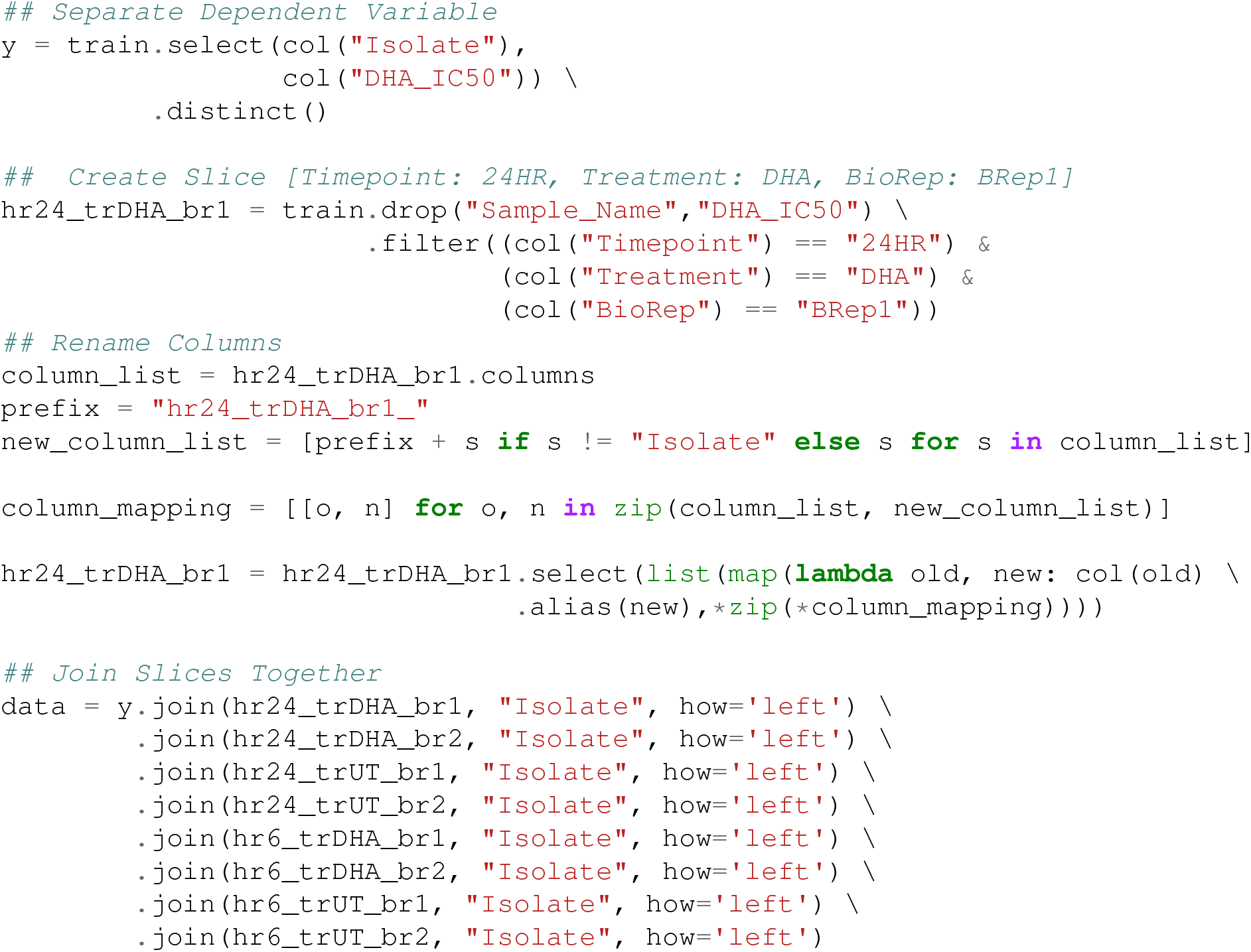

Lastly, the dataset is then vectorized using the Spark `VectorAssembler`, and converted into a Numpy^12^-compatible array. Example code shown below in Code 1. Vectorization allows for highly scalable parallelization of the machine learning modeling in the next step.

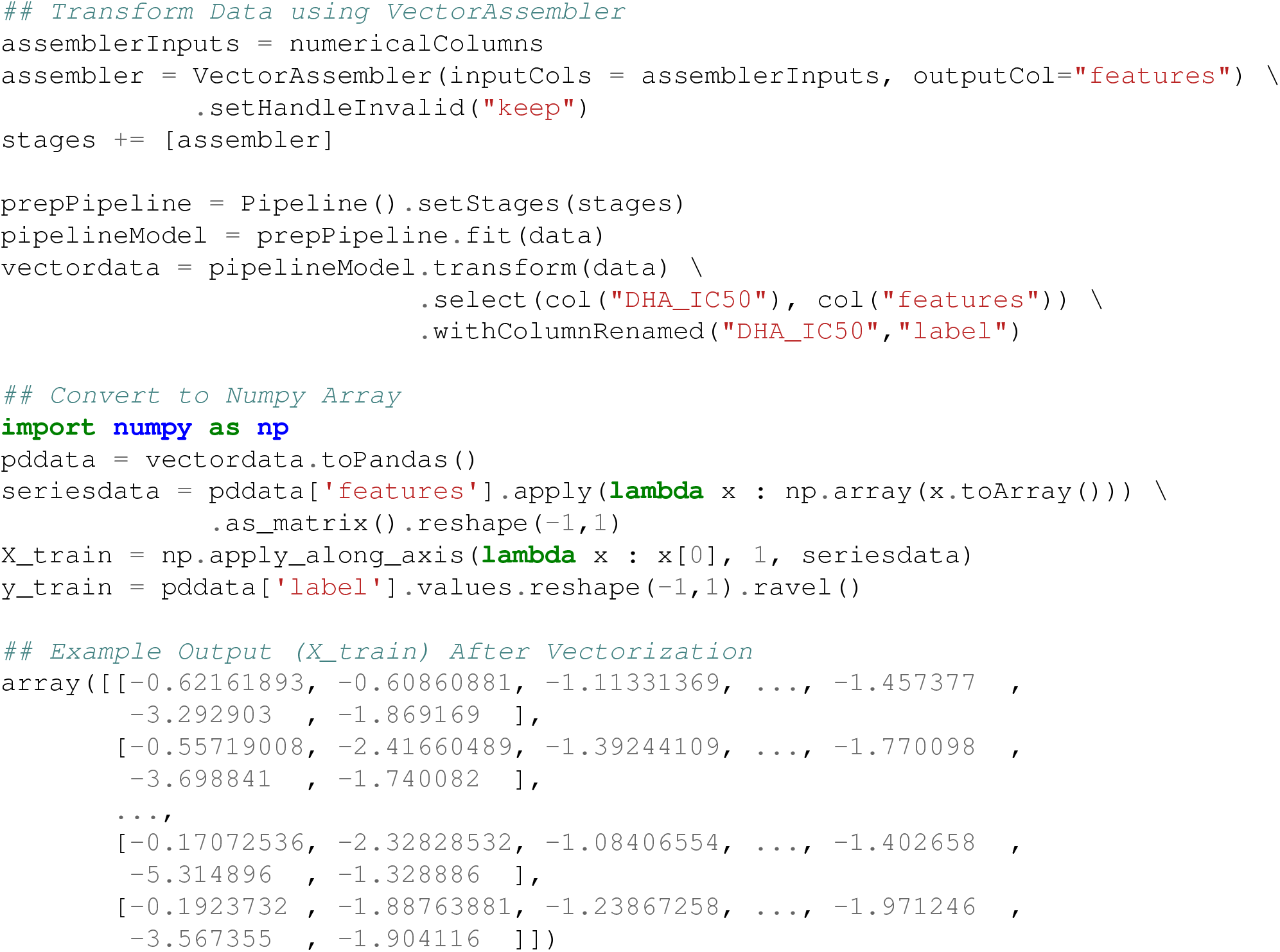

### Machine Learning

We used Microsoft Azure Machine Learning Service as the tracking platform for retaining model performance metrics as the various models were generated. For this use case, 498 machine learning models were trained using various scaling techniques and algorithms. We then created two ensemble models of the individual models using Stack Ensemble and Voting ensemble methods.

The Microsoft AutoML package allows for the parallel creation and testing of various models, fitting based on a primary metric. For this use case, models were trained using Decision Tree, Elastic Net, Extreme Random Tree, Gradient Boosting, Lasso Lars, LightGBM, RandomForest, and Stochastic Gradient Decent algorithms along with various scaling methods from Maximum Absolute Scaler, Min/Max Scaler, Principal Component Analysis, Robust Scaler, Sparse Normalizer, Standard Scale Wrapper, Truncated Singular Value Decomposition Wrapper (as defined in Table 14). All of the machine learning algorithms are from the *scikit-learn* package^13^ except for LightGBM, which is from the *LightGBM* package^14^. That the settings for the model sweep are defined in Table 4. The ‘Preprocess Data?’ parameter enables the scaling and imputation of the features in the data.

**Table 4.** Model search parameter setting for the IC_50_ model search.

| Parameter | Value |
| --- | --- |
| Task | Regression |
| Number of Iterations | 500 |
| Iteration Timeout (minutes) | 20 |
| Max Cores per Iteration | 7 |
| Primary Metric | Normalized Root Mean Squared Error |
| Preprocess Data? | True |
| k-Fold Cross-Validations | 20 folds |

Once the 498 individual models were trained, two ensemble models (voting ensemble and stack ensemble) were then created and tested. The voting ensemble method makes a prediction based on the weighted average of the previous models’ predicted regression outputs whereas the stacking ensemble method combines the previous models and trains a meta-model using the elastic net algorithm based on the output from the previous models. The model selection method used was the Caruana ensemble selection algorithm^15^.

## Results

The voting ensemble model (using soft voting) was selected as the best model, having the lowest normalized Root Mean Squared Error (RMSE), as shown in Table 5. All 500 models trained are reported in Table 6. Having a normalized RMSE of only 0.1228 and a Mean Absolute Percentage Error (MAPE) of 24.27%, this model is expected to accurately predict IC_50_ in malaria isolates.

**Figure 1.**
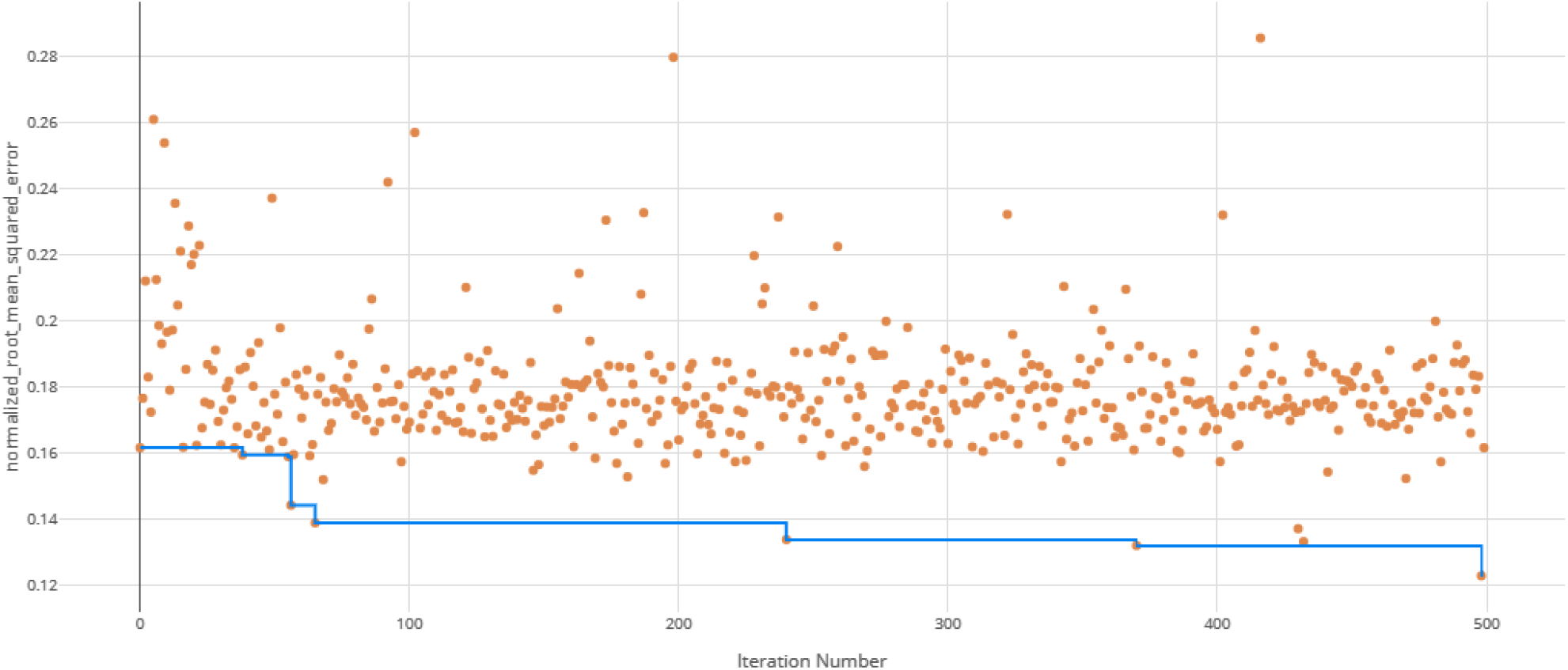
RMSE by iteration of the IC_50_ model search. Each orange dot is an iteration with the blue line representing the minimum RMSE up to that iteration.

**Figure 2.**
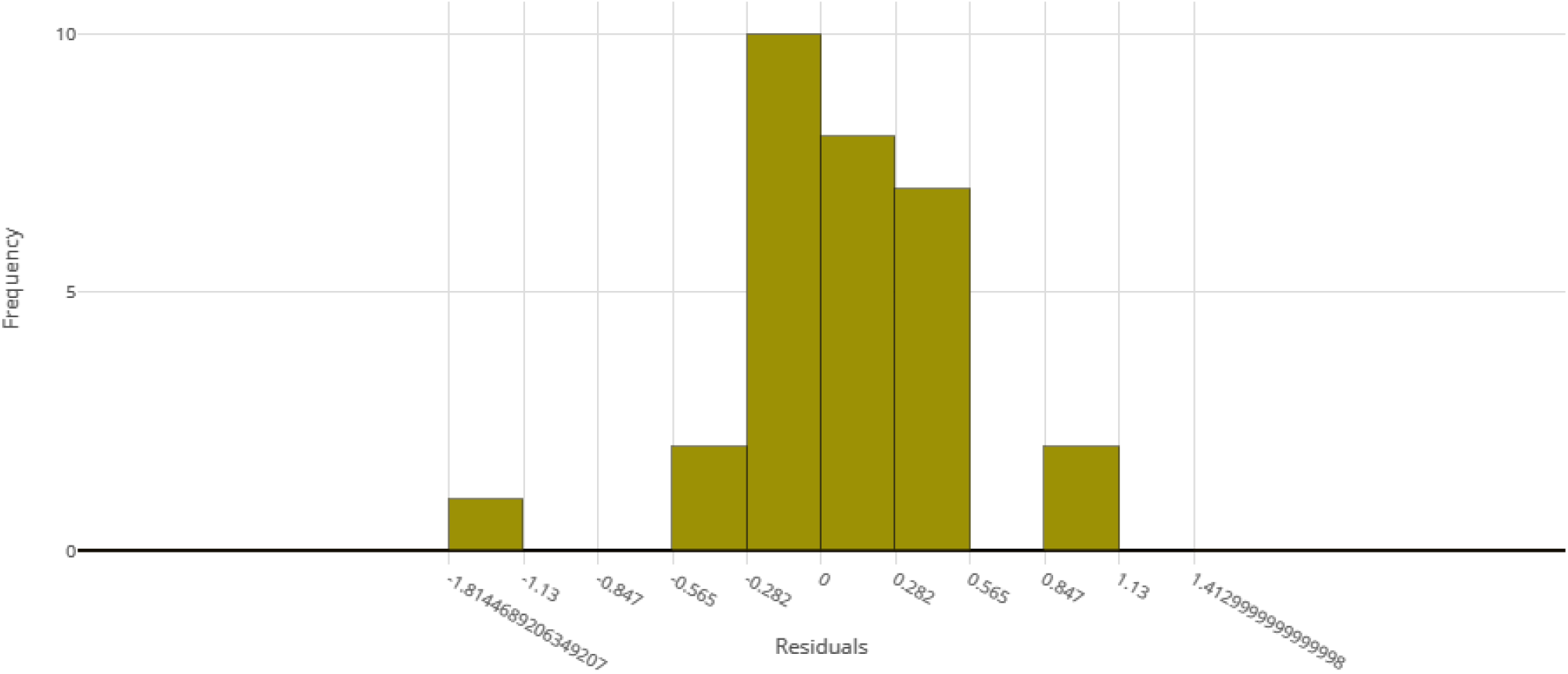
Model residuals of the final IC_50_ ensemble model.

**Table 5.** Model metrics of the final IC_50_ ensemble model.

| Metric | Value |
| --- | --- |
| Normalized Root Mean Squared Error | 0.1228 |
| Root Mean Squared Log Error | 0.1336 |
| Normalized Mean Absolute Error | 0.1097 |
| Mean Absolute Percentage Error | 24.27 |
| Normalized Median Absolute Error | 0.1097 |
| Root Mean Squared Error | 0.3398 |
| Explained Variance | -1.755 |
| Normalized Root Mean Squared Log Error | 0.1379 |
| Median Absolute Error | 0.3035 |
| Mean Absolute Error | 0.3035 |

**Table 6.**
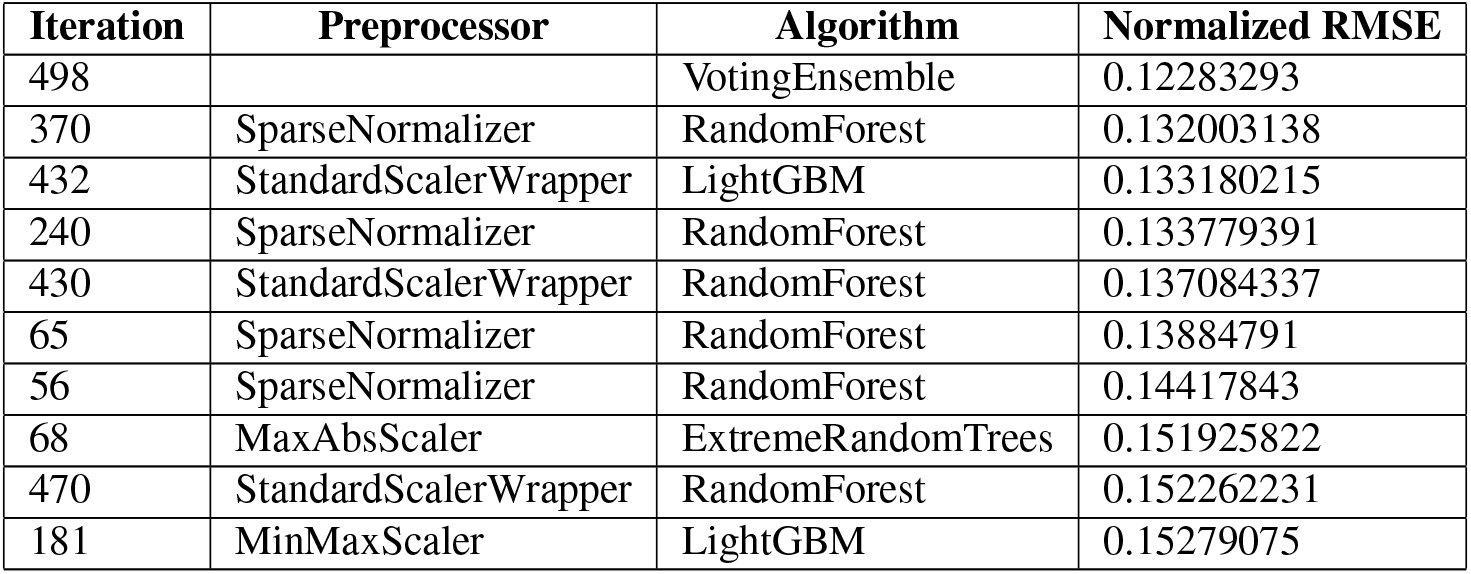
Top 10 training iterations of the IC_50_ model search, evaluated by RMSE. Note that the top performing model (VotingEnsemble) is the final IC_50_ model discussed in this paper.

| Iteration | Preprocessor | Algorithm | Normalized RMSE |
| --- | --- | --- | --- |
| 498 |  | VotingEnsemble | 0.12283293 |
| 370 | SparseNormalizer | RandomForest | 0.132003138 |
| 432 | StandardScalerWrapper | LightGBM | 0.133180215 |
| 240 | SparseNormalizer | RandomForest | 0.133779391 |
| 430 | StandardScalerWrapper | RandomForest | 0.137084337 |
| 65 | SparseNormalizer | RandomForest | 0.13884791 |
| 56 | SparseNormalizer | RandomForest | 0.14417843 |
| 68 | MaxAbsScaler | ExtremeRandomTrees | 0.151925822 |
| 470 | StandardScalerWrapper | RandomForest | 0.152262231 |
| 181 | MinMaxScaler | LightGBM | 0.15279075 |

## Prediction of Resistance Status

The second task of this work was to create a machine learning model that can predict the parasite clearance rate (fast versus slow) of malaria isolates. When resistance rates change in a pathogen, it can be indicative of regulatory changes in the pathogen’s genome. These changes can be exploited for the prevention of further resistance spread. Thus, a goal of this work is to understand genes important in the prediction of artemisinin resistance.

### Materials and Methods

An *in vivo* transcription data set from Mok et al., (2015) Science^16^ was used to predict the parasite clearance rate of malaria parasite isolates based on *in vitro* transcriptional profiles. See Table 8.

**Table 7.** Format of the clearance rate model training data.

| Sample_Names | Country | Asexual_stage__hpi__ | Kmeans_Grp | PF3D7_0100100 | ... | PF3D7_1480100 | ClearanceRate |
| --- | --- | --- | --- | --- | --- | --- | --- |
| GSM1427365 | Bangladesh | 20 | B | 0.226311 | ... | -0.64171 | Fast |
| ... | ... | ... | ... | ... | ... | ... | ... |
| GSM1427537 | Cambodia | 12 | C | 0.81096 | ... | -1.72825 | Slow |
| ... | ... | ... | ... | ... | ... | ... | ... |
| GSM1428407 | Vietnam | 8 | A | 0.999095 | ... | NaN | Fast |

**Table 8.** Training dataset information from Mok et al., 2015^16^.

|  | Training Set |
| --- | --- |
| Number of Isolates | 1043 |
| Isolate Collection Site | Southeast Asia |
| Isolate Collection Years | 2012-2014 |
| Sample Type | *in vivo* |
| Synchronized? | Not Synchronized |
| Number of Samples per Isolate | 1 |
| Additional Attributes | ~18 hpi,<br>Non-perturbed,<br>No replicates |

The training data consists of 1,043 isolates with 4,952 genes from the malaria parasite, *Plasmodium falciparum*. For each malaria parasite isolate, transcription data was collected for various *PF3D7* genes. The form of the training dataset contains 1,043 rows and 4,957 columns, as shown in Table 7. The feature to predict is *ClearanceRate*.

### Data Preparation

The training data for this use case did not require the same pivoting transformations as in the last use case as each record describes a single isolate. Thus, only the vectorization of the data was necessary, which was performed using the Spark `VectorAssembler` and then converted into a Numpy-compatible array^12^. Example code shown below in Code 1. Note that this vectorization only kept the numerical columns, which excludes the `Country, Kmeans_Grp`, and `Asexual_stage__hpi_` attributes as they are either absent or contain non-matching factors (i.e. different set of countries) in the testing data.

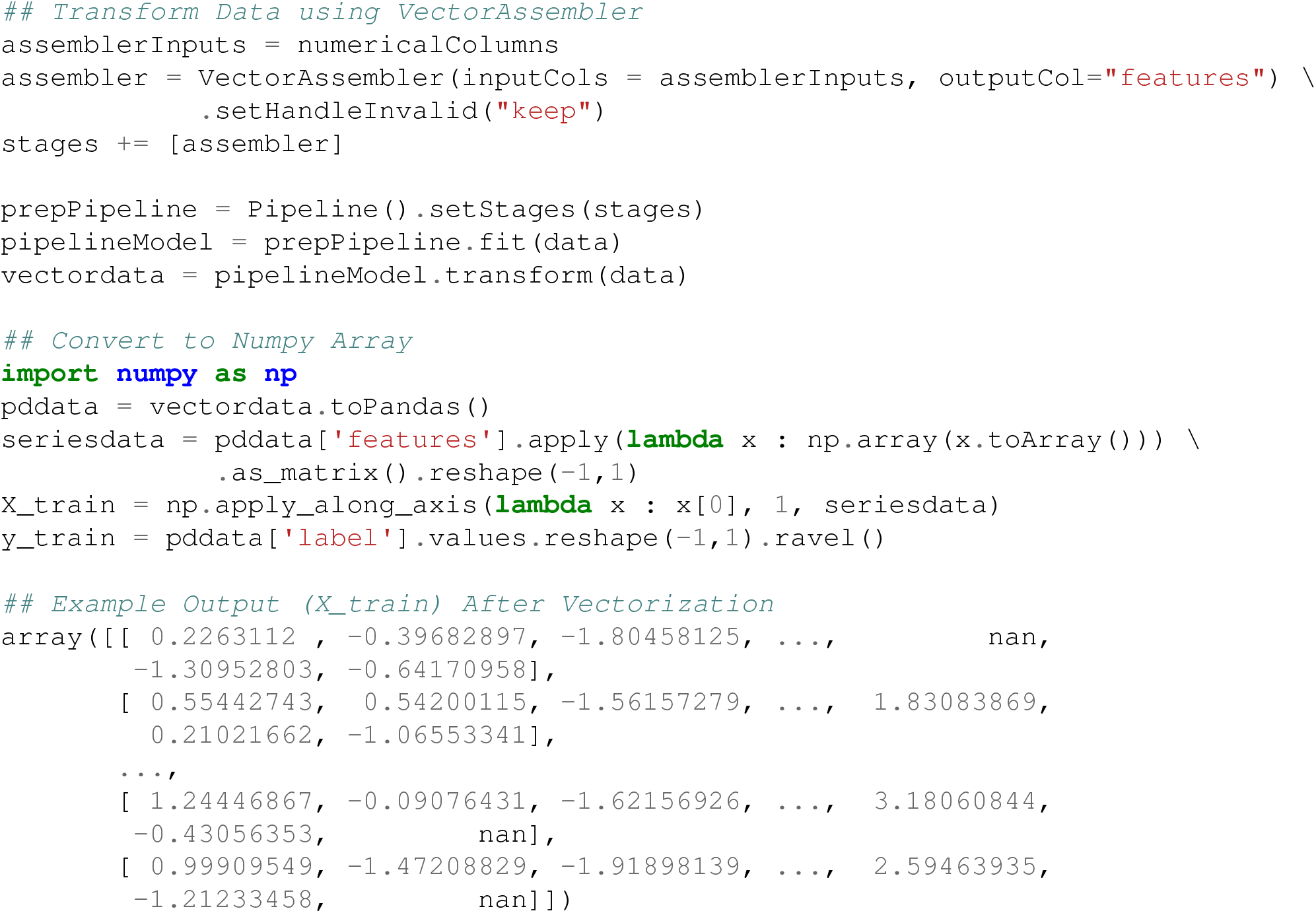

### Machine Learning

Once the 98 individual models were trained, two ensemble models (voting ensemble and stack ensemble) were then created and tested as before.

## Results

The voting ensemble model (using soft voting) was selected as the best model, having the highest Area Under the Receiver Operating Characteristic curve (AUC), as shown in Table 11. The top 10 of the 100 models trained are reported in Table 10. Having a weighted AUC of 0.87 and a weighted F1 score of 0.80, this model is expected to accurately predict isolate clearance rates.

**Table 9.** Model search parameter settings for the clearance rate model search.

| Parameter | Value |
| --- | --- |
| Task | Regression |
| Number of Iterations | 100 |
| Iteration Timeout (minutes) | 20 |
| Max Cores per Iteration | 14 |
| Primary Metric | Weighted AUC |
| Preprocess Data? | True |
| k-Fold Cross-Validations | 10 folds |

**Table 10.** Top 10 training iterations of the clearance rate model search. Note that the top performing model (VotingEnsemble) is the clearance rate model discussed in this paper.

| Iteration | Preprocessor | Algorithm | Weighted AUC |
| --- | --- | --- | --- |
| 98 |  | VotingEnsemble | 0.870471056 |
| 99 |  | StackEnsemble | 0.865215516 |
| 65 | StandardScalerWrapper | LogisticRegression | 0.86062304 |
| 33 | StandardScalerWrapper | LogisticRegression | 0.859881677 |
| 97 | StandardScalerWrapper | LogisticRegression | 0.858791006 |
| 44 | StandardScalerWrapper | LogisticRegression | 0.856105491 |
| 73 | StandardScalerWrapper | LogisticRegression | 0.855502817 |
| 17 | RobustScaler | SVM | 0.855452622 |
| 43 | StandardScalerWrapper | LogisticRegression | 0.855368394 |
| 61 | RobustScaler | LogisticRegression | 0.854357599 |

**Table 11.**
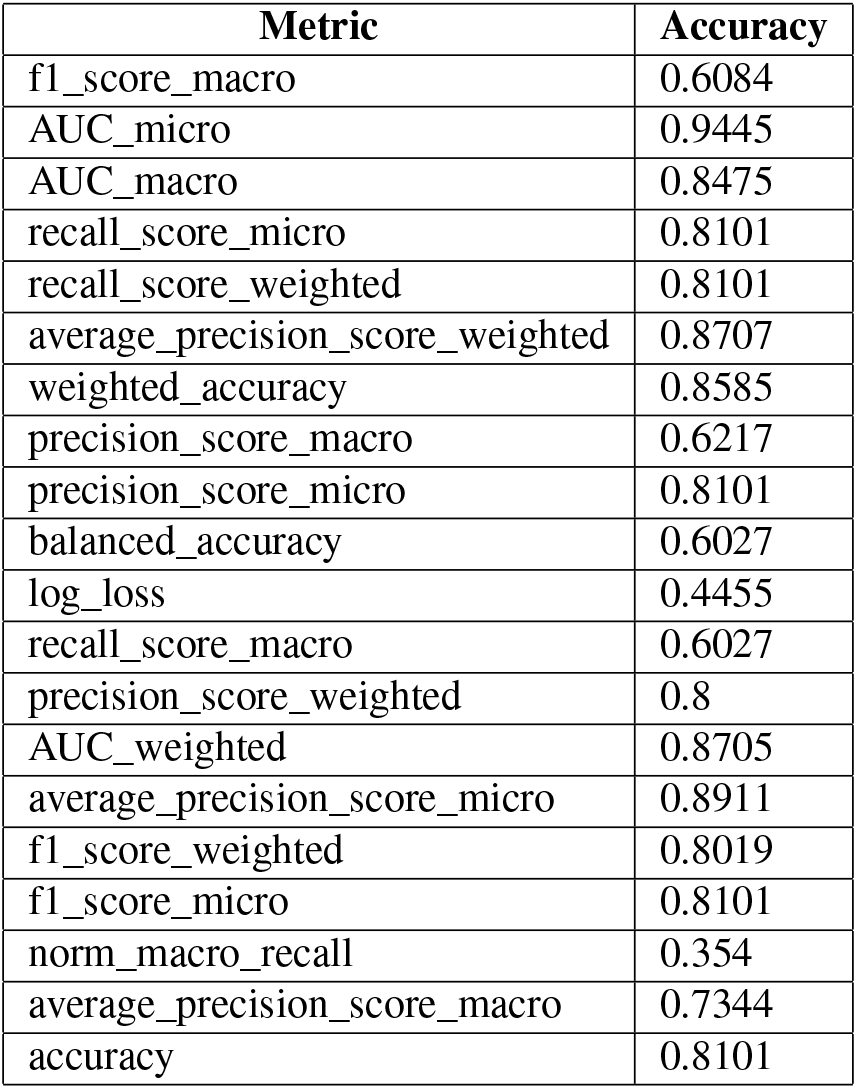
Model metrics of the final clearance rate ensemble model.

| Metric | Accuracy |
| --- | --- |
| f1_score_macro | 0.6084 |
| AUC_micro | 0.9445 |
| AUC_macro | 0.8475 |
| recall_score_micro | 0.8101 |
| recall_score_weighted | 0.8101 |
| average_precision_score_weighted | 0.8707 |
| weighted_accuracy | 0.8585 |
| precision_score_macro | 0.6217 |
| precision_score_micro | 0.8101 |
| balanced_accuracy | 0.6027 |
| log_loss | 0.4455 |
| recall_score_macro | 0.6027 |
| precision_score_weighted | 0.8 |
| AUC_weighted | 0.8705 |
| average_precision_score_micro | 0.8911 |
| f1_score_weighted | 0.8019 |
| f1_score_micro | 0.8101 |
| norm_macro_recall | 0.354 |
| average_precision_score_macro | 0.7344 |
| accuracy | 0.8101 |

### 0.1 Feature Importance

Feature importances were calculated using mimic-based model explanation of the ensemble model. The mimic explainer works by training global surrogate models to mimic blackbox models. The surrogate model is an interpretable model, trained to approximate the predictions of a black box model as accurately as possible. See Figure 6 and Table 13.

**Figure 3.**
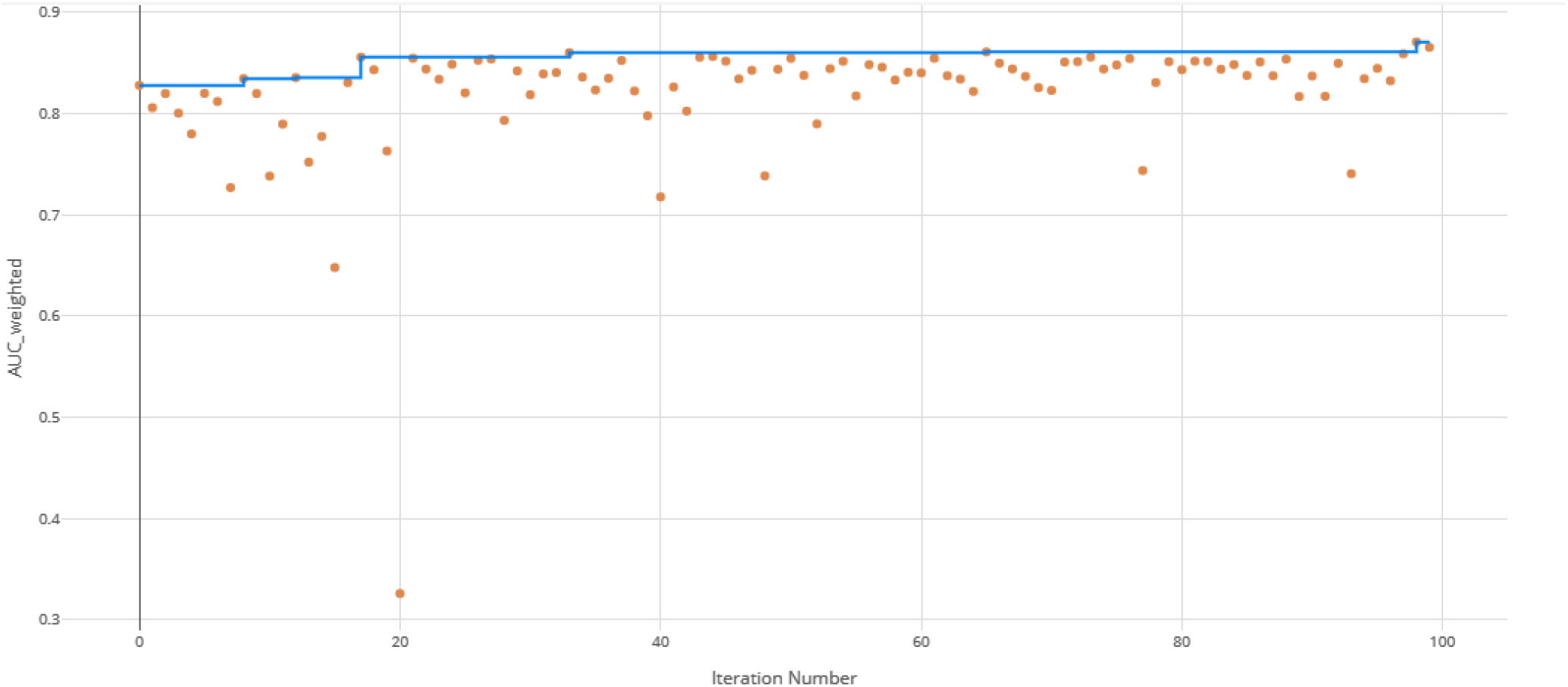
AUC by iteration of the clearance rate model. Each orange dot is an iteration with the blue line representing the maximum AUC up to that iteration.

**Figure 4.**
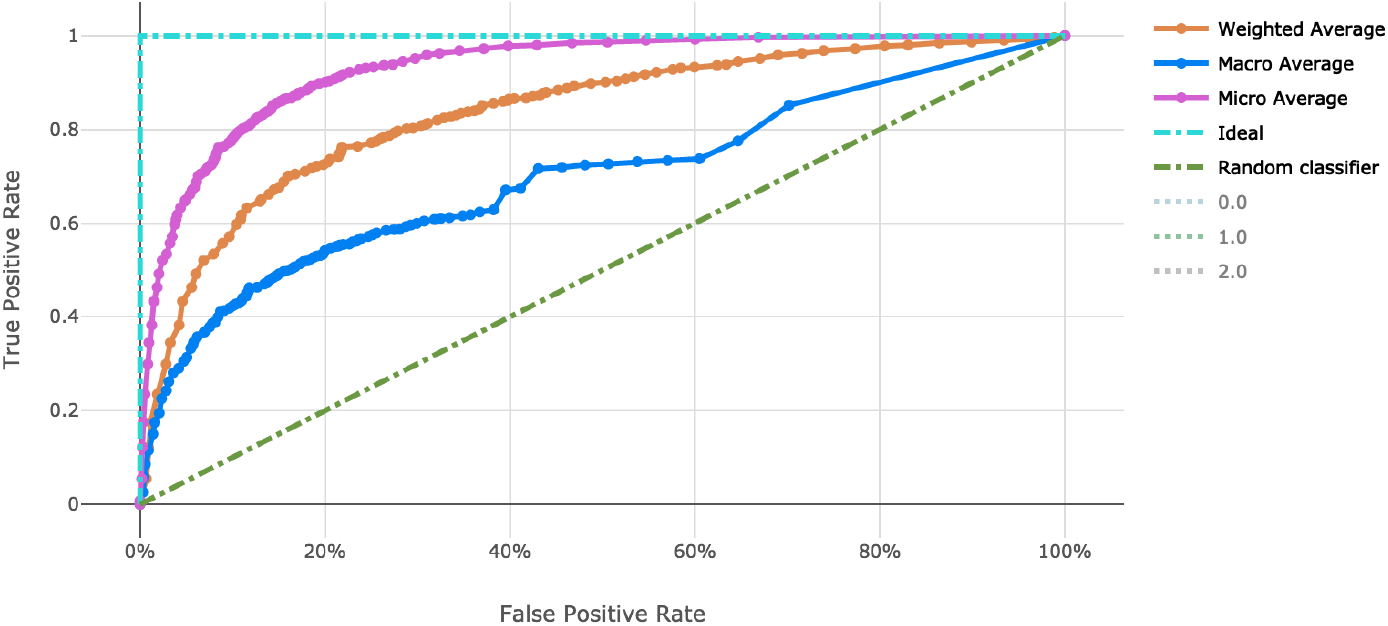
ROC curve of the clearance rate model.

**Figure 5.**
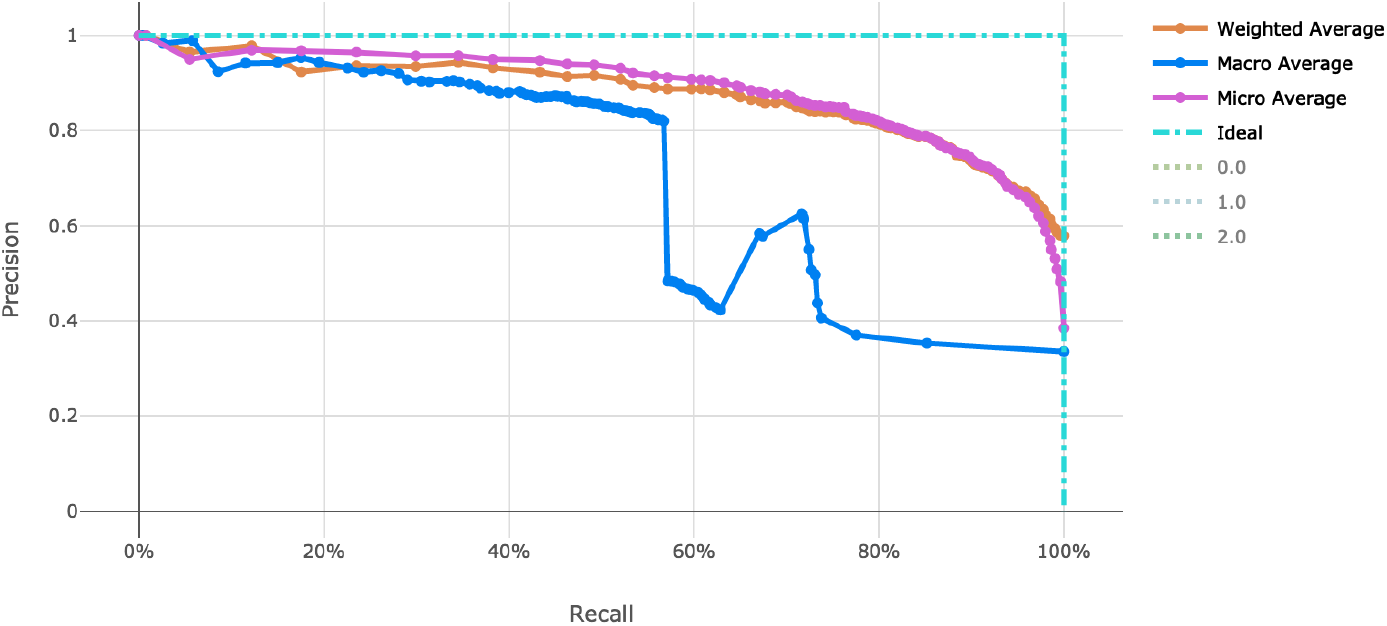
Precision-Recall curve of the clearance rate model.

**Figure 6.**
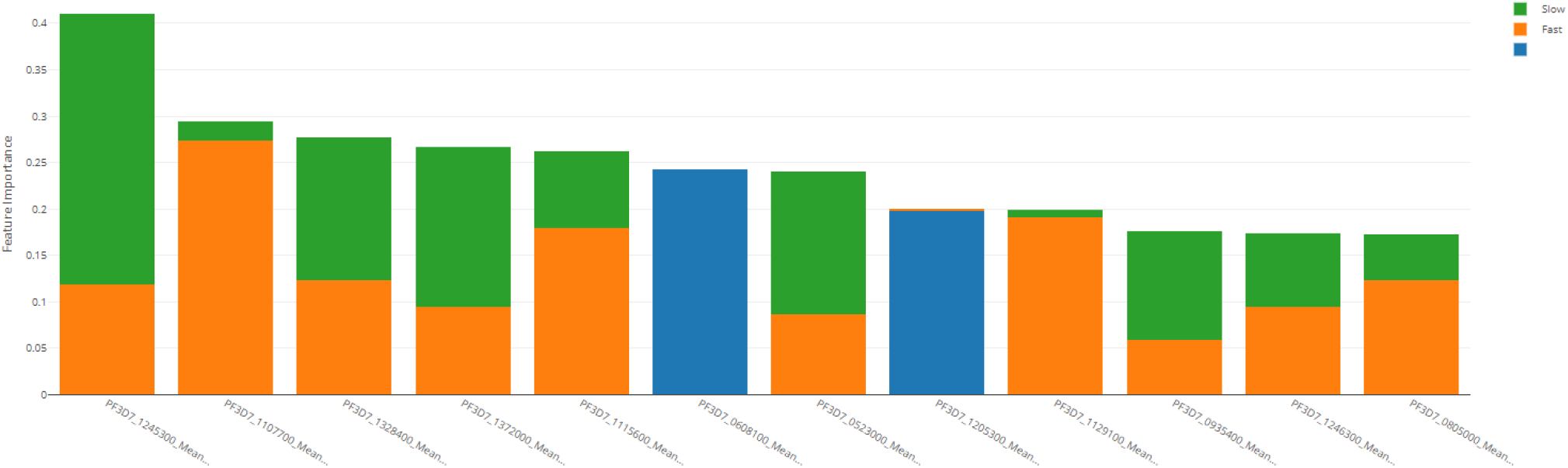
Derived feature importances using the black box mimic model explanation of the clearance rate model

**Table 13.** Top 10 PF3D7 genes (features) in predicting clearance rate.

| Rank | PF3D7 Gene | "Fast" Importance | "Slow" Importance | Overall Importance |
| --- | --- | --- | --- | --- |
| 1 | 1245300 | 0.292 | 0.118 | 0.41 |
| 2 | 1107700 | 0.02 | 0.274 | 0.294 |
| 3 | 1328400 | 0.154 | 0.123 | 0.277 |
| 4 | 1372000 | 0.172 | 0.095 | 0.267 |
| 5 | 1115600 | 0.083 | 0.179 | 0.262 |
| 6 | 0523000 | 0.154 | 0.087 | 0.241 |
| 7 | 1129100 | 0.008 | 0.191 | 0.199 |
| 8 | 0935400 | 0.117 | 0.058 | 0.175 |
| 9 | 1246300 | 0.079 | 0.095 | 0.174 |
| 10 | 0805000 | 0.049 | 0.124 | 0.173 |

**Table 12.** Confusion matrix of clearance rate predictions versus actual.

|  |  | Prediction |  |  |
| --- | --- | --- | --- | --- |
|  |  | Fast (ID: 0) | Slow (ID: 1) | Null (ID: 2) |
| Actual | Fast (ID: 0) | 661 | 74 | 0 |
|  | Slow (ID: 1) | 115 | 184 | 0 |
|  | Null (ID: 2) | 6 | 3 | 0 |

## Discussion

By using distributed processing of the data preparation, we can successfully shape and manage large malaria datasets. We efficiently transformed a matrix of over 40,000 genetic attributes for the *IC*_50_ use case and over 4,000 genetic attributes for the resistance rate use case. This was completed with scalable vectorization of the training data, which allowed for many machine learning models to be generated. By tracking the individual performance results of each machine learning model, we can determine which model is most useful. In addition, ensemble modeling of the various singular models proved effective for both tasks in this work.

The resulting model performance of both the *IC*_50_ model and the clearance rate model show relatively adequate fitting of the data for their respective predictions. While additional model tuning may provide a lift in model performance, we have demonstrated the utility of ensemble modeling in these predictive use cases in malaria.

In addition, this exercise helps to quantify the importance of genetic features, spotlighting potential genes that are significant in artemisinin resistance. The utility of these models will help in directing development of alternative treatment or coordination of combination therapies in resistant infections.

## Supplementary Materials

**Table 14.** Scaling function information for machine learning model search.

| Scaling and Normalization | Description |
| --- | --- |
| StandardScaleWrapper | Standardize features by removing the mean and scaling to unit variance |
| MinMaxScaler | Transforms features by scaling each feature by that column's minimum and maximum |
| MaxAbsScaler | Scale each feature by its maximum absolute value |
| RobustScaler | This Scaler features by their quantile range |
| PCA | Linear dimensionality reduction using singular value decomposition of the data to project it to a lower dimensional space |
| TruncatedSVDWrapper | This transformer performs linear dimensionality reduction by means of truncated singular value decomposition.<br>Contrary to PCA, this estimator does not center the data before computing the singular value decomposition. This means it can efficiently work with sparse matrices. |
| SparseNormalizer | Each sample (each record of the data) with at least one non-zero component is re-scaled independently of other samples so that its norm (L1 or L2) equals one |

All code, datasets, models, and results are hosted at http://github.com/colbyford/malaria_DREAM2019.

## Acknowledgements

The Datasets used for the analyses described in this manuscript were contributed by Michael T. Ferdig, Department of Biological Sciences at the University of Notre Dame. They were obtained as part of the Malaria DREAM Challenge through Synapse (syn16924919), managed by Sage Bionetworks.

## Author contributions statement

C.T.F. conceived and conducted the data transformation and machine learning modeling, C.T.F. and D.J. analysed the results. Both authors reviewed the manuscript.

## Additional information

The authors declare no competing interests.

